# Impaired Functional Connectivity of Cortico-Amygdala Pathway Can Drive Social Behavior Deficits in Synucleinopathies

**DOI:** 10.1101/2024.05.20.594995

**Authors:** Wei Zhou, Samuel Daniels, Vijay Singh, Marissa Menard, Martha L Escobar Galvis, Hong-Yuan Chu

## Abstract

The small molecule protein α-synuclein forms insoluble aggregates in a group of neurological disorders, including Parkinson’s disease and dementia with Lewy bodies (DLB), which are collectively called synucleinopathies. In PD and DLB, the amygdala has been identified as a particularly susceptible region in the brain for the deposition of Lewy-like α-synuclein aggregates. Though α-synuclein aggregation is closely associated with neurodegeneration, there is a poor correlation between neurodegeneration in the amygdala and the clinical features of PD/DLB. We hypothesize that, prior to neurodegeneration, α-synuclein aggregation disrupts functional cortical modulation of the amygdala circuits, leading to emotion dysregulation in synucleinopathies. In the present study, we combined electrophysiology, optogenetics, mouse model of synucleinopathies, and behavioral analysis to test this hypothesis. Using an α-synuclein preformed fibrils (PFFs)-based mouse model of synucleinopathies, we reported dynamic changes in the levels of α-synuclein pathology in the basolateral amygdala (BLA). Such dynamic changes of pathology associated with a decreased cortico-BLA connection strength prior to a significant loss of cortical axon terminals. In parallel to the reduced cortico-BLA connection, PFFs-injected mice manifested impaired social preference behavior. The impaired sociability of PFFs-injected mice could be rescued by chemogenetic stimulation of cortico-BLA inputs. Altogether, we presented a series of evidence to delineate key circuit events associated with α-synuclein pathology development in the amygdala circuits. The present work highlights the necessity of a thorough investigation of functional consequences of α-synuclein aggregation to advance our understand of pathophysiology of synucleinopathies and development of effective therapies.

## Background

Abnormally folded and aggregated α-synuclein (α-Syn) is the major proteinaceous component of the intraneuronal inclusions that characterize Lewy pathology [1, 2]. Lewy pathology can be found in most brains of people with Parkinson’s disease (PD), dementia with Lewy bodies (DLB), and multiple system atrophy. These disorders are collectively known as “synucleinopathies” [2]. Experimental work suggests that gradual accumulation of α-Syn aggregates can underlie the pathogenesis and progression of synucleinopathies [3–5]. Particularly, accumulation of pathologic α-Syn aggregates has been reported to impair nigrostriatal dopaminergic neurotransmission and closely relate to the neurodegeneration, leading to the manifestation of motor symptoms in PD [4, 6, 7]. In addition, α-Syn pathology is proposed to spread between cells and brain regions [3, 5, 8, 9]. Consequently, a large body of studies have focused on understanding the genetic and microenvironmental factors that may promote or prevent the formation of pathologic α-Syn aggregation, as they perhaps can provide us with strategies to regulate the temporal development and spatial distribution of α-Syn pathology across brain regions and in that way modify disease progression [10–15].

Formation of α-Syn aggregates in the brain trigger series of circuitry events, including but not limited to, the degeneration of vulnerable neuronal populations. For example, compelling evidence also suggests that α-Syn aggregates disrupt synaptic structure, function, and plasticity in susceptible brain regions, like the nigrostriatal pathway [4, 16–18]. Motor dysfunction in PD is closely linked with dopamine depletion in the basal ganglia and the subsequent functional maladaptation within the basal ganglia-thalamocortical network [19, 20]. Similar pathophysiological processes are likely to occur in other brain regions that are vulnerable to the formation of α-synuclein aggregates, in which functional circuit changes, not necessarily neurodegeneration, are causally linked to symptom manifestation. The amygdala has been reported as a preferential brain region for Lewy pathology development in PD and DLB [21–24] and animal models of synucleinopathy [3, 25, 26].

Considering the critical role of the amygdala in emotion regulation, heavy load of Lewy-like pathology in the amygdala and the subsequent neuronal loss are expected to causally associate with depression and/or anxiety in PD/DLB [27–30].However, such a linear relationship between neurodegeneration and clinical features has not been supported by studies using human tissues of PD/DLB [24] or its animal models [25, 31]. Building on the research about the nigrostriatal dopaminergic system, it is plausible to hypothesize that impaired amygdala circuit function, but not significant neurodegeneration, is sufficient to cause emotion dysregulation in synucleinopathies. In the present work, we combined electrophysiological, immunohistochemical, and behavioral approaches to test this hypothesis. Using an intrastriatal α-synuclein preformed fibrils (PFFs) model, we demonstrated that: 1) α-synuclein aggregates selectively eliminate cortical afferents to the basolateral amygdala (BLA); 2) α-synuclein aggregation induces functional impairment of cortico-BLA inputs, prior to robust synapse loss; 3) neuronal loss in the BLA occurs at late stages as α-synuclein aggregates accumulate; and 4) social behavioral deficits associate with cortico-BLA synaptic impairments, occurring prior to synaptic or neuronal loss in the BLA, and can be alleviated by functional restoration of cortico-BLA connection strength. Our work here provides novel insights into the amygdala circuit dysfunction associated with α-Syn pathology formation and possible biological mechanisms underlying emotion dysregulation in PD and DLB.

## Materials and methods

### Animals

C57BL/6J mice at 2-3-month-old (Jax#:000664, RRID: IMSR_JAX:000664) of both sexes were obtained through the Van Andel Institute vivarium and used in this study. All animal studies were reviewed and approved by the Institutional Animal Care and Use Committee at Van Andel Institute (animal use protocol #: 22-02-007) and in accordance with the standards of NIH for care and use of animals. Animals were housed under a 12:12 h light-dark cycle, up to four animals per cage with access to water and food *ad libitum*.

### Preparation and validation of α-Syn preformed fibrils

The mouse α-Syn protein was purified using *Escherichia coli* BL21 codon plus RIPL cells (RRID:CVCL_M639), which was then dialyzed using a buffer containing 10 mM Tris and 50 mM NaCl (pH 7.5). A high-capacity endotoxin removal kit (PI88276) was used to remove the endotoxins, the levels of which were assessed using an endotoxin quantification kit (A39552). The protein concentration was estimated using absorbance at 280 nm with an extinction coefficient of 7450 M^−1^ cm^−1^. Purified mouse α-Syn monomer protein was used to generate mouse α-Syn preformed fibrils (PFFs). Specifically, monomeric α-Syn protein was diluted to 5 mg/mL in the buffer, 150 mM KCl, 50 mM Tris-HCl and incubated at 37°C with shaking for 7 days, as described previously [32]. After the incubation, sample was centrifuged for 10 min at 13,200 rpm. The protein pellet was re-suspended in half of the initial volume of the solution. To estimate the fibril concentration, 5 μl of fibril solution was incubated with 5 μL of 8 M guanidinium chloride at room temperature for one hour. After incubation, the concentration of PFF was measured using absorbance at 280 nm and diluted the PFF at 5 mg/mL and 22-25 μl aliquots were stored at −80°C until use. Prior to the injection, PFF aliquot (22-25 μl at 5 mg/mL) was thawed at room temperature and sonicated using Qsonica 700W cup horn sonicator at 30% amplitude using 3 seconds on/2 seconds off cycle for 15 min at 15°C. The size of sonicated PFF (30-70 nm segments) was estimated and confirmed using the dynamic light scattering (DynaPro NanoStar from Wyatt technology). Detailed of generating and validating α-Syn PFFs can be found here: dx.doi.org/10.17504/protocols.io.bhhrj356.

### Stereotaxic surgery

Motorized stereotaxic frames were used for stereotaxic injections (Model# 51730M, Stoelting Co.). Mice were anesthetized with 2% isoflurane and placed on a heating pad throughout the surgery for maintenance of the body temperature. Eye ointment was administered to keep eyes moisture and to protect them from strong light. To induce α-Syn pathology in mouse brain, we bilaterally injected the sonicated PFFs (10 μg in a 2.0 μL volume) into the dorsal striatum (anteroposterior (A-P), +0.2 mm from bregma; mediolateral (M-L), ± 2.0 mm from the midline; dorsoventral (D-V), −2.9 mm from the brain surface) using a Nanoliter injector (NANOLITER2020, World Precision Instrument) at a rate of 0.4 μL/min. To label the BLA principal neurons for electrophysiological recordings, Retrobeads (LumaFluor Inc) were diluted (1:10 dilution) into PFFs or PBS and then the mixed materials were co-injected into the dorsal striatum. Mice injected with PBS or PBS plus beads at the same age were made as control groups. To optogenetically evoke excitatory postsynaptic currents (EPSCs) of BLA projection neurons, we injected 0.3 μL AAVs encoding ChR2(H134R)-eYFP (titer = 1 x 10^12^ GC/ml, Addgene# 26973, RRID: Addgene_127090) into the layer 5 of medial prefrontal cortex (mPFC, A-P, +2.1 mm from bregma; M-L, ± 0.3 mm from the midline; D-V, −2.3 mm from the brain surface) and the centromedial thalamus (A-P, −1.5 mm from bregma; M-L, 0 mm from the middle; D-V, −3.3 mm from the brain surface) at a rate of 0.1 μL/min. An intersectional approach was used for selective activation of mPFC to BLA projection using chemogenetics: 1) retrograde AAV encoding FlpO (0.3 μL, titer ≥ 7 x 10^12^ GC/ml, Addgene# 55637; RRID:Addgene_55637) were bilaterally injected into BLA (A-P, −1.3 mm from bregma; M-L, ± 3.3 mm from the middle; D-V, −4.4 mm from the brain surface), and 2) AAVs encoding FlpO-dependent excitatory designer receptors exclusively activated by designer drugs (DREADDs) (titer = 2.5 x 10^13^ GC/ml, Addgene 154868; RRID:Addgene_154868) were injected into infralimbic subregion of mPFC (IL-mPFC) (A-P, +2.1 mm from bregma; M-L, ± 0.3 mm from the midline; D-V, −2.6 mm from the brain surface).

Animals were returned to a cage placed on a warm pad after surgery and then housed in their home cage until experiments. Details of this protocol can be found at: dx.doi.org/10.17504/protocols.io.rm7vzye28lx1/v1

### Slice preparation for *ex vivo* physiological recording

To prepare brain slices for physiological recording, mice were intraperitoneally injected with avertin (300 mg/kg) for anesthesia, followed by transcardial perfusion using ice-cold, sucrose-based artificial cerebrospinal fluid (aCSF), containing the follows (in mM): 230 sucrose, 26 NaHCO_3_, 10 glucose, 10 MgSO_4_, 2.5 KCl, 1.25 NaH_2_PO_4_, 0.5 CaCl_2_, 1 sodium pyruvate, and 0.005 L-glutathione. Brain slices (300 μm thickness) with the BLA were collected using a vibratome (VT1200S; Leica Microsystems, Buffalo Grove, IL) in the same sucrose-aCSF. Throughout the preparation, brain slices were kept at about +4°C by a recirculating chiller (FL300; JULABO USA, Inc, Allentown, PA). Then the slices were incubated at 35°C in regular aCSF bubbled with 95% O_2_ and 5% CO_2_ for 30 min (in mM, 126 NaCl, 26 NaHCO_3_, 10 glucose, 2.5 KCl, 2 CaCl_2_, 2 MgSO_4_, 1.25 NaH_2_PO_4_, 1 sodium pyruvate, and 0.005 L-glutathione), and then kept at room temperature for at least another 30 min prior to recording. Details of this procedure can be found here: https://www.protocols.io/view/protocol-for-obtaining-rodent-brain-slices-for-ele-ewov1y7mpvr2/v2

### Slice electrophysiology recording and optogenetics

Brain slices were transferred to recording chamber with modified aCSF (in mM, 126 NaCl, 26 NaHCO3,10 glucose, 3 KCl, 1.6 CaCl_2_,1.5 MgSO_4_, and 1.25 NaH_2_PO_4_), which was equilibrated with 95% O_2_ and 5% CO_2_ at 32–34°C via a feedback-controlled in-line heater (TC-324C, Warner Instruments). A gradient contrast SliceScope 1000 (Scientifica, UK) with an IR-2000 CCD camera (DAGE-MTI, USA) using infrared illumination was used to visualize neurons in slices. A motorized micromanipulator was used to control the position of the glass pipette for recording. The BLA principal cells were identified based on the presence of somatic Retrobeads signals under a 60x water immersion object lens (Olympus, Japan).

Electrophysiology data were collected using an Axon 700B amplifier and digitized with a 1550B digitizer (Molecular Devices, San Jose, USA). Signals were low pass filtered at 2 kHz and sampled at 50 kHz under the control of pClamp 11 (RRID: SCR_011323, Molecular Devices, San Jose, USA). Borosilicate glass pipettes with resistance at 4-6 MΩ (O.D. = 1.5 mm, I.D. = 0.86, item # BF150-86-10, Sutter Instruments, Novato, CA) were made by a micropipette puller (P1000, Sutter Instruments, Novato, CA). For voltage-clamp recordings of synaptic transmission, a Cs^+^-based intracellular solution containing the following (in mM): 120 CH_3_O_3_SCs, 2.8 NaCl, 10 HEPES, 0.4 Na_4_-EGTA, 5 QX314-HBr, 5 phosphocreatine, 0.1 spermine, 4 ATP-Mg, and 0.4 GTP-Na was used; the pH was adjusted to pH 7.3 with CsOH. A K-gluconate-based internal solution (in mM, 140 K-gluconate, 3.8 NaCl, 1 MgCl2, 10 HEPES, 0.1 Na_4_-EGTA, 2 ATP-Mg, and 0.1 GTP-Na, 0.2% Biocytin, pH 7.3, 290 mOsm) was used to label the BLA projections neurons for morphology studies using Cy-streptavidin. For optogenetic stimulation, light pulses with 1 ms duration were delivered through a 60x objective lens (Olympus, Japan) using a 470 nm LED light illumination (CoolLED, UK). TTX (1 μM) and 4-AP (100 μM) were perfused to isolate monosynaptic mPFC-BLA neurotransmission. To measure the synaptic strength, peak amplitude of AMPARs-mediated EPSCs at −80 mV were calculated. Paired-pulse ratio at 10 Hz was quantified as the ratio of amplitude between the second EPSCs over the first EPSCs (i.e., EPSC2/EPSC1). To measure AMPA/NMA ratio, NMDA-mediated current at +40 mV was measured at 50 ms after optogenetic stimulation when AMPAR mediated-EPSCs decayed to the baseline. Series resistance (R_s_) was continuously monitored during recordings and only cells showed stable R_s_ < 20 MΩ with less than 15% changes were proceeded for further analysis. Details of this procedure can be found here: https://www.protocols.io/view/brain-slice-preparation-for-electrophysiology-reco-36wgqj2eovk5/v1

### Perfusion and immunohistochemistry staining

Mice were intraperitoneally injected with avertin for anesthesia followed by transcardial perfusion using 1x PBS and 4% paraformaldehyde (PFA, pH = 7.4). After perfusion, the brains were extracted and kept in 4% PFA overnight for post-fixation, and then were re-sectioned at a 70 μm thickness using a vibratome (VT1000s, Leica Biosystems, Buffalo Grove, IL). https://www.protocols.io/view/standard-operating-procedure-mouse-transcardiac-pe-3byl4b8r8vo5/v1

Brain sections were rinsed with 1x PBS for 3 times before performing immunohistochemistry, followed by incubation with 1x PBS containing 0.5% Triton X-100 (PBS-T) plus 2% normal donkey serum (Millipore Sigma) at room temperature for 60 min. They were then incubated in the PBS-T with primary antibodies overnight at room temperature or 48 hours at 4°C. The primary antibodies utilized in this study were: rabbit anti-pS129 α-Syn (1:10,000, Bioscience Cat# 1536-1, RRID: AB_562180), mouse anti-NeuN (1:500, Millipore Cat# MAB377, RRID: AB_2298772), mouse anti-vGluT1 (1:1,000, Synaptic Systems Cat# 135 011, RRID:AB_2617087), guinea pig anti-vGluT2 (1:1,000, Synaptic System Cat# 135404, RRID: AB_887884). The sections were then thoroughly rinsed with 1x PBS for 3 times before incubating with the secondary antibodies (1: 500, AlexaFluor 594 donkey anti-rabbit IgG, Cat# 711-585-152, Jackson ImmunoResearch Labs, RRID: AB_2340621; AlexFluor 594 donkey anti-mouse IgG, Cat# 715-586-150, Jackson ImmunoResearch Labs, RRID: AB_2340857; AlexaFluor 488 donkey anti-guinea pig IgG, Cat# 706-545-148, Jackson ImmunoResearch Labs, RRID: AB_2340472) at room temperature for 2 hours. To reveal morphology of biocytin-labeled neurons, brain slices were incubated with Cy5-conjugated streptavidin (SA1011, Thermo Fisher Scientific) at a concentration of 1: 500 at room temperature for 2 h. Brain slices were then rinsed using 1x PBS for 3 times before mounting on glass slides with the Vectorshield antifade mounting medium (H-1000, Vector Laboratories). Slides were left at room temperature overnight before sealing with nail oil and imaging. Details of immunofluorescent staining can be found here: https://www.protocols.io/view/immunofluorescent-staining-3byl4bq9ovo5/v1

### Confocal imaging and analysis

Confocal images were taken using a Nikon A1R microscope. Images of α-Syn pathology in BLA were collected by a 20x objective lens and then analyzed using imageJ (RRID: SCR_003070, NIH, https://imagej.net). To quantify the cortical and thalamic axon terminals in the BLA, vGluT1 and vGluT2 immunoreactivity were imaged using a Nikon A1R confocal microscope under an oil immersion 100x objective lens (NA=1.45; x/y, 1024 x 1024 pixels; z-step = 150 nm). All confocal images were acquired using the identical microscope and camera settings. VGluT1 and vGluT2 density were stereologically quantified using optical dissector methods (dissector height = 3 μm with a guard zone of 0.15 μm; counting frame = 10 x 10 μm) [33]. Images of NeuN staining were taken from the BLA by a 60x objective (NA=1.40; x/y, 1024 x 1024 pixels; z-step = 1 μm) and subjected to quantification using Imaris software (RRID: SCR_007370, version 10.0, Oxford, UK, https://imaris.oxinst.com). ChR2-eYFP expression in mPFC was imaged using a Nikon Ni-U fluorescence microscope under 2x objective lens. For morphology analysis, biocytin-filled neurons were imaged using the confocal microscope under 20x objective lens (NA=0.75; x/y, 1024 x 1024 pixels; z-step = 1 μm). Morphology of biocytin-labelled neurons of BLA was reconstructed followed by Sholl analysis using the Imaris software (RRID: SCR_007370, version 10.0, Oxford, UK, https://imaris.oxinst.com). Spines of the biocytin-filled neurons were imaged by the confocal microscope under 100x objective lens (NA=1.45; x/y, 1024 x 1024 pixels; z-step = 500 nm). To quantify the spine density of BLA neurons, dendritic segments of 20 μm in length were reconstructed and number of spines were counted. Images from controls and PFFs injected mice were collected under identical settings in the confocal microscope and analyzed by a researcher blinded to treatments. Details of digital imaging analysis can be found here: https://www.protocols.io/view/confocal-imaging-and-digital-image-analysis-3byl4jmxzlo5/v1.

### Animal behaviors

To assess motor function, animals were subject to locomotion and rotarods tests. For open field locomotion test, the animal was placed in the empty open field (40 x 40 x 30 cm, W x L x H) and allowed to explore for 10 min. Animal’s locomotor activity was continuously recorded and analyzed using ANYmaze software (RRID:SCR_014289, Stoelting Co., Wood Dale, IL; http://www.sandiegoinstruments.com/any-maze-video-tracking/). For rotarod test, the animal was placed on an accelerating rod (speed of 4-40 rpm in 5 min, San Diego Instruments, CA) and the latency of fall was monitored and recorded. Detailed protocol of motor activity assays can be found here: https://www.protocols.io/view/open-field-locomotion-test-e6nvwjxmdlmk/v1. To assess animals’ social interaction behavior, mice were placed to three-chamber apparatus (MazeEngineers, Skokie, IL) with left, middle, and right chambers (20 x 13.5 cm^2^). A round wire cage (10 cm in diameter, 20 cm in height) was placed in the left and right chambers for novel stimulus mouse. The stimulus mice were habituated to the wire cages for 20 min before the testing day. The testing mice were habituated to the behavior room for at least 30 min before the experiment. Three sessions were conducted during the testing day, including the habituation, the session 1, and the session 2. The testing mice were habituated to the middle chamber for 5 min before opening the doors to allow the mice freely explore the three chambers for 10 min. During the first session, the testing mice were allowed to explore the three chambers for a total of 10 min with one empty cage and the presence of a stimulus mouse in the other wire cage. In the second session, the same testing mouse was allowed to explore the three chambers for another 10 min with the presence of the familiar mouse in one cage and a second never-met stimulus mouse in another wire cage. Mice movements were tracking and the time spent in each chamber was quantified by AnyMaze software (RRID:SCR_014289, Stoelting Co.). Between each testing, the chamber was thoroughly cleaned with 70% ethanol to eliminate the odor of mice.

### Data analysis and statistics

Electrophysiology data were analyzed using Clampfit 11.1 (RRID: SCR_011323, Molecular Devices, San Jose, USA). The amplitude of monosynaptic glutamatergic EPSCs were quantified from an average of three sweeps. Confocal images were analyzed using Image J (RRID: SCR_003070, NIH, https://imagej.net) and Imaris (RRID: SCR_007370, Oxford, UK, http://www.bitplane.com/imaris). Statistics were performed in Prism 10 (RRID: SCR_002798, GraphPad Software, http://www.graphpad.com). Non-parametric, distribution-independent Mann-Whiney U (MWU) test was used for statistical comparison of data from two groups. The amplitude of Cortico-BLA EPSCs across a range of stimulation and Sholl analysis were analyzed using two-way analysis of variance (ANOVA) with *post hoc* Sidak’s analysis multiple comparison’s test. All tests were two-tailed, and *p*-value < 0.05 was considered as statistical significance.

## Results

### Time-course of α-Syn pathology development in the BLA

We injected α-Syn PFFs into the dorsal striatum of wildtype C57BL6 mice to induce the accumulation of α-Syn pathology in the brain, including the cerebral cortical regions and the amygdala [3, 25, 31]. In the PFFs-based models, fibrillar α-Syn species are reported to be internalized at the axon terminals, which then recruit endogenous α-Syn to form insoluble inclusions and eventually result in neurodegeneration [3, 11, 16].

As expected, we detected robust phospho-Ser129 (pS129^+^) α-Syn immunoreactivity in the BLA of PFFs-injected mice (Figure 1A-D), but not in the BLA of PBS- or monomer-injected controls (“controls” hereinafter). α-Syn pathology in the BLA included both pS129^+^ Lewy neurite (LN)-like and large Lewy body (LB)-like aggregates (Figure 1A-D), which are likely the α-Syn pathology within cortical afferents of the BLA and the cytoplasmic aggregates within glutamatergic principal neurons of the BLA, respectively [31, 34]. Next, we characterized the dynamic changes of α-Syn pathology abundance in the BLA at 1, 3, 6, and 12 months-post-injection (mpi). We measured the proportion of the BLA region covered by LN- and LB-like pS129^+^ aggregation, as well as their combination as the total α-Syn pathology (i.e., pS129^+^ LN/LB-like aggregates, Figure 1E) [25, 26]. Specifically, moderate level of pS129^+^ α-Syn pathology was detected in the BLA as early as at 1 mpi, which was dominated by the LN-like pS129^+^ aggregates (Figure 1A**, E**). The level of total pS129^+^ α-Syn pathology continually grew beyond 1 mpi, peaking at 3 mpi (Figure 1B**, E**). After 3 mpi, the amount of total α-Syn pathology in the BLA remained at a relatively high at the 6 mpi timepoint but decreased significantly at 12 mpi (Figure 1C**, D, E**). Particularly, at 12 mpi only few LB-like pS129^+^ aggregates could be seen in the BLA (Figure 1D**, E**).

**Figure 1.**
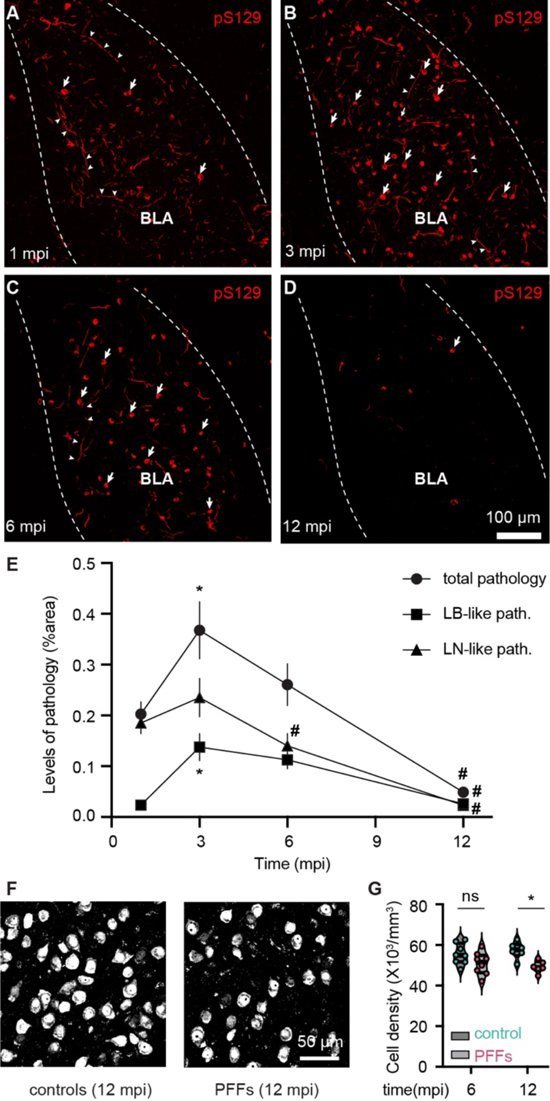
Time-course of αSyn pathology development in the BLA. **A-D)** Representative confocal images showing the development and accumulation of pS129^+^ pathology in the BLA at 1, 3, 6 and 12 months-post-injection (mpi). Arrows and arrowheads highlighted a few pS129^+^ somatic aggregates and neurites, respectively, in the BLA. **E)** Time course of the development of total α-syn pathology (filled circles), LB-like pS129^+^ pathology (square), and LN-like pS129^+^ pathology (triangle) in the BLA. Total α-Syn pathology, 1 mpi = 0.20±0.02, n = 38 slices/8 mice; 3 mpi = 0.37±0.06, n = 28 slices/10 mice; p = 0.004. LB-like pS129^+^ pathology, 1 mpi =0.02±0.005, n = 38 slices/8 mice; 3 mpi =0.14±0.03, n = 28 slices/10 mice, p < 0.0001. LN-like pS129^+^ pathology, 1 mpi =0.19±0.02, n = 38 slices/8 mice, 3 mpi = 0.24±0.04, n = 28 slices/10 mice, p = 0.3. Total α-Syn pathology at 6 mpi = 0.26±0.04, n = 38 slices/10 mice, p = 0.09 versus total pathology at 3 mpi. LB-like pS129^+^ pathology at 6 mpi = 0.11±0.02, n = 38 slices/10 mice, p = 0.5 versus LB-like pS129^+^ pathology at 3 mpi. LN-like pS129^+^ pathology at 6 mpi = 0.14±0.02, n = 38 slices/10 mice, p = 0.01 versus LN-like pS129^+^ pathology at 3 mpi. Total α-Syn pathology at 12 mpi = 0.05±0.007, n = 35 slices/6 mice, p < 0.0001 versus total pathology at 3 mpi. LB-like pS129^+^ pathology at 12 mpi = 0.026±0.005, n = 35 slices/6 mice, p < 0.0001 versus LB-like pS129^+^ pathology at 3 mpi. LN-like pS129^+^ pathology at 12 mpi = 0.02±0.0035, n = 35 slices/6 mice, p < 0.0001 versus LN-like pS129^+^ pathology at 3 mpi. *p< 0.05 relative to the counterparts at 1 mpi, #p < 0.05 relative to the counterparts at 3 mpi; one-way ANOVA. **F**) Representative images showing NeuN immunoreactivity in the BLA of controls and PFFs-injected mice at 12 mpi. **G**) Summarized results showing a reduced BLA cell density in the PFFs-injected mice relative to controls at 12 mpi, but not at the 6 mpi. Cell densities at 6 mpi, control = 54.9 [51.7, 60.8] x10^3^/mm^3^, PFFs = 51.5 [46.4, 54.9] x10^3^/mm^3^, n = 8 mice/group, p = 0.16; Cell densities at 12 mpi, control = 57.7 [54.7, 60.3] x10^3^/mm^3^, PFFs = 48.9 [47.4, 51.8] x10^3^/mm^3^, n = 6 mice/group, p = 0.0087, Mann-Whitney U (MWU) test.

Post-mortem studies of patients with advanced PD reported that the amygdala exhibits significant Lewy body pathology and reduced cell density [24]. Consistently, using unbiased stereological counting of pan-neuronal marker NeuN, we detected a significant reduction of the NeuN density in the BLA at 12 mpi relative to controls, but not at 6 mpi (Figure 1F**, G**, see also [31]). These data are consistent with the loss of cortical glutamatergic neurons and midbrain dopaminergic neurons driven by the formation and maturation of LB-like pS129^+^ aggregates [3, 11, 35]. Together the above results suggest that robust reduction in the LB-like pS129^+^ aggregates at 12 mpi was, at least partially, due to neuronal loss in the BLA.

### Loss of cortical afferents contributes to the decreased pS129^+^ neurites in the BLA

Glutamatergic inputs from the cerebral cortex and the thalamus are the major excitatory drive of the BLA neuronal activity and they form synapses with the spines and dendrites of BLA principal neurons. Interaction of cortical and thalamic inputs is critical to proper manifestation of emotion and behavior. Our recent study suggest that cortical and thalamic axons show distinct functional vulnerability to α-Syn aggregation at 1 month post PFFs injections when there was no overt loss of glutamatergic axonal terminals. It remains to be determined whether and when α-Syn aggregates induce loss of cortical and/or thalamic inputs in the BLA. Thus, we rigorously examined cortical and thalamic inputs to the BLA at additional time points in both controls and PFFs-injected mice.

Cortical and thalamic axon terminals can be distinguished by their preferential expression of vesicular glutamate transporter (vGluT) 1 and 2, respectively. Using optical dissector methods, we stereologically quantified the density of vGluT1^+^ puncta in the BLA, starting from 3 mpi when the level of LN-like pS129^+^ α-Syn aggregation reached a peak (Figure 1E). Although pS129^+^ α-Syn was tremendous in the BLA at 3 mpi, there was no change in the density of vGluT1^+^ puncta (Figure 2A). However, a significant reduction in the density of vGluT1^+^ puncta in the BLA was detected at 6 mpi (Figure 2B), indicating a gradual loss of cortico-BLA axonal projections at later time points, though a downregulation of vGluT1 protein itself cannot be excluded.

**Figure 2.**
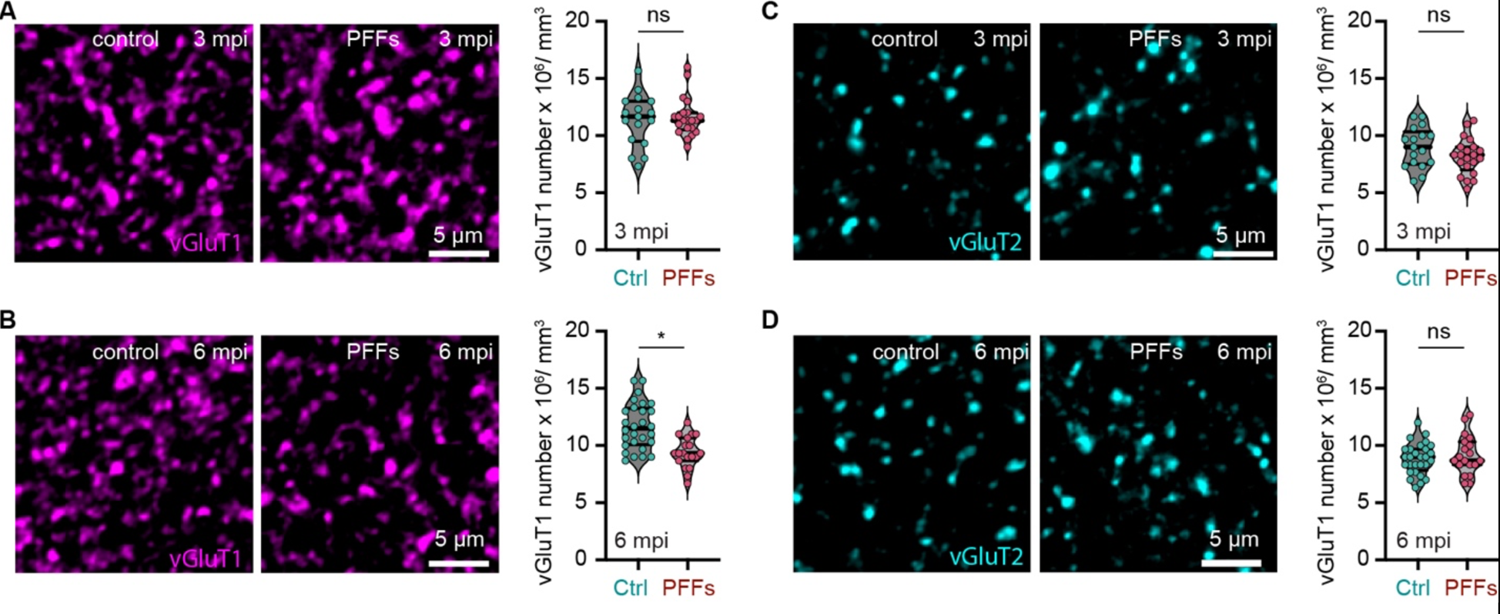
Loss of cortical afferents contributes to the decreased pS129^+^ neurites in the BLA. **A-B**) *Left*, representative confocal images showing vGluT1 immunoreactive puncta in the BLA of controls and PFFs-injected mice at 3 and 6 mpi. *Right*) Summarized result showed changes in vGluT1 densities in the BLA at 3 and 6 mpi. *A,* vGluT1 densities at 3 mpi, controls = 11.7 [9.5, 13], PFFs = 11.3 [10.5, 12.0], p = 0.86. *B,* vGluT1 densities at 6 mpi, controls = 11.5 [10.1, 13.3], PFFs = 9.3 [8.7, 10.7], p = 0.0001. **C-D**) *Left*, representative confocal images showing vGluT2 immunoreactive puncta in the BLA of controls and PFFs-injected mice at 3 and 6 mpi. *Right*) Summarized result showed no change in vGluT2 densities in the BLA at 3 and 6 mpi. *D,* vGluT2 densities at 3 mpi, controls = 9.0 [7.5, 10.3], PFFs = 8.3 [7.0, 9.0], p = 0.1387. *E,* vGluT2 densities at 6 mpi, controls = 9.0 [7.8, 9.7], PFFs = 8.7 [8.3, 10.3], p = 0.4883. N = 21 slices/4 mice for each group at 3 mpi, 19 slices/4 mice for each group at 6 mpi. MWU test.

When it came to thalamic inputs to the BLA, we found no change in the density of vGluT2^+^ puncta in the BLA of PFFs injected mice relative to respective controls at 3 and 6 mpi (Figure 2C**, D**). These data indicate that thalamic-BLA axonal projections were not altered by the α-Syn pathology in the brain. Taken together, we concluded that the reduction of LN-like pS129^+^ aggregates in the BLA between 3 mpi and 6 mpi was mainly due to a loss of cortical axonal inputs, but not the thalamic afferents.

Given the cortical inputs mainly target dendritic spines of BLA principal neurons, loss of cortical axonal terminals is expected to be accompanied by a reduction of postsynaptic spines.

Thus, we further quantified the density of dendritic spines of the biocytin labeled BLA principal neurons that were retrogradely labeled by intrastriatal injection of Retrobeads and PFFs mixture (Figure 3A**, B**). We found that BLA principal neurons exhibited a reduced density of dendritic spines at 3 and 6 mpi (Figure 3C-F).

**Figure 3.**
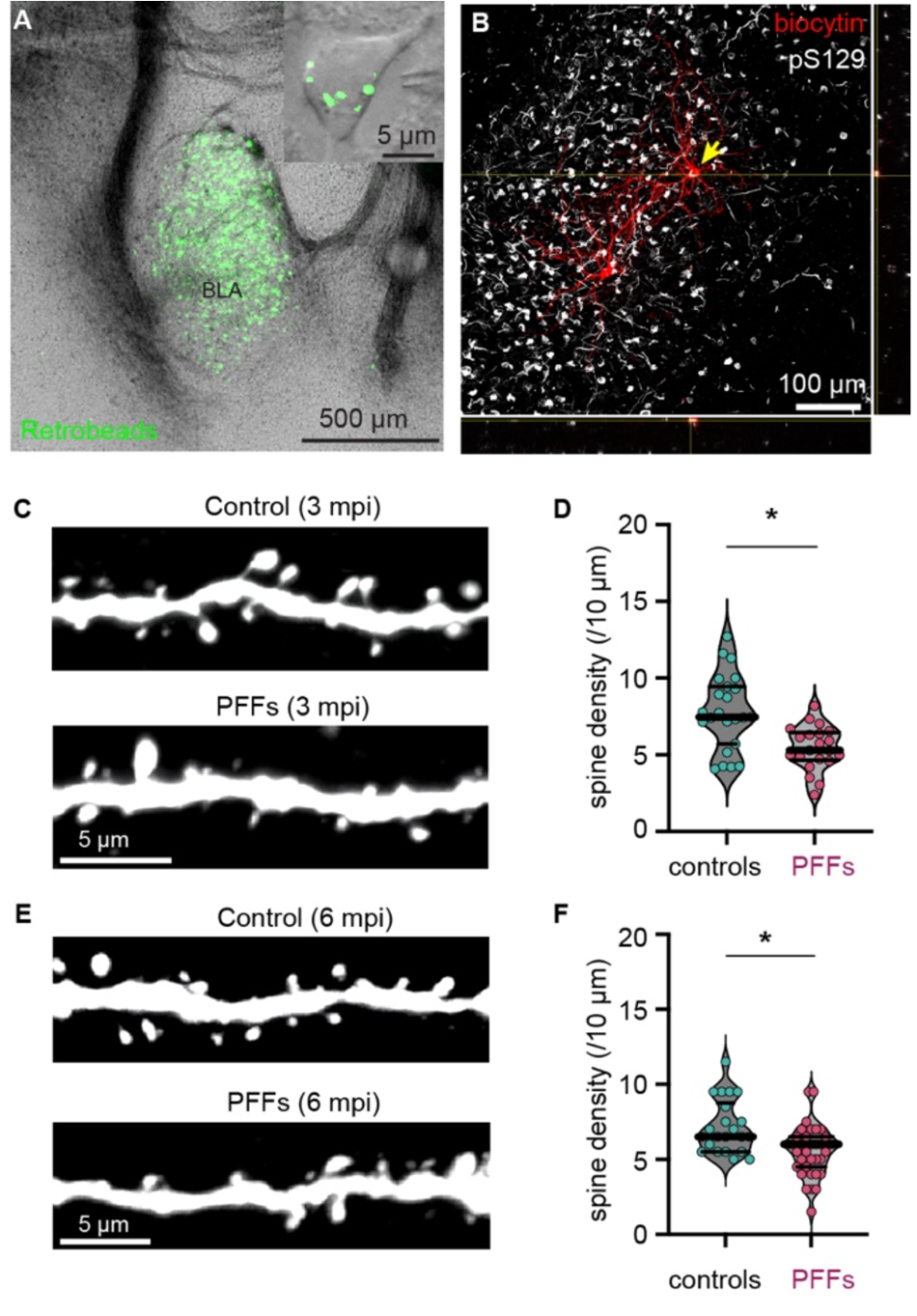
Loss of dendritic spines of the BLA principal neurons as α-Syn pathology develops. **A)** Representative images showing green Retrobeads-labeled BLA neurons at low and high (inset) magnification. Cells with green beads labeling were targeted for biocytin filling and morphology analysis. **B**) Representative image showing a Cy5-labeled BLA principal neurons against pS129^+^ α-Syn immunoreactivity. This cell showed clear somatic aggregation of pS129^+^ α-Syn. **C**) Confocal images showing representative dendritic spines of BLA neurons from controls and PFFs-injected mice at 3 mpi. **D**) Summarized results showing reduced dendritic spine densities of BLA neurons from PFFs-injected mice relative to control at 3 mpi. **E-F**) Similar to (C-D) but at 6 mpi. Spine densities at 3 mpi, controls = 7.45 [5.7, 9.44]/10 μm; PFFs = 5.3 [4.66, 6.46]/10 μm, n= 21 segments/4 mice, p = 0.0008, MWU; Spine densities at 6 mpi, controls = 6.5 [5.5, 8.75]/10 μm, PFFs = 6.0 [4.5, 6.5]/10 μm, n = 36 segments/4 mice, p = 0.002, MWU.

Taken together, α-Syn aggregation triggers loss of presynaptic cortical axon terminals and postsynaptic dendritic spines of BLA principal neurons, which is likely to result in a loss of cortico-BLA synapses as α-Syn pathology accumulates in the brain, e.g. at 6 mpi.

### α-Syn pathology disrupts cortico-BLA functional connectivity prior to loss of synapses

After establishing a time course of neuronal and synaptic losses in the BLA, we next assessed the functional connectivity between the cerebral cortex and the BLA. The medial prefrontal cortex (mFPC) and the BLA form interconnected microcircuits that play a critical role in regulating social behavior [36–38]. Both the mPFC and the BLA accumulate robust α-Syn pathology following intrastriatal PFFs injection[3, 31], the functional connectivity of mPFC-BLA pathway is expected to be impaired. Thus, we assessed how α-Syn pathology affects mFPC-BLA connection at 3 and 6 mpi when the abundance of α-Syn pathology remained high but has caused no or subtle degeneration of BLA neurons and presynaptic cortical axonal terminals (Figures 1 **and 2**). We virally expressed AAV9-hSyn-ChR2(H134R)-eYFP in the layer 5 projection neurons of the mPFC, allowing a selective activation of mPFC-BLA inputs (Figure 4A). We optogenetically stimulated ChR2(H134R)-expressing axon terminals in the BLA (Figure 4A) and recorded the evoked excitatory postsynaptic currents (EPSCs) in the retrogradely labeled BLA principal neurons under the voltage-clamp mode from controls and PFFs-injected mice (Figure 4B**, C**). The amplitude of optogenetically-evoked, monosynaptic EPSCs were quantified to assess the connection strength of mPFC-BLA synapses. A significant reduction of the amplitude of mPFC-BLA EPSCs was detected in PFFs-injected mice relative to control at both 3 and 6 mpi (Figure 4B**, C**), indicating a reduced connection strength of the mPFC-BLA synapses.

**Figure 4.**
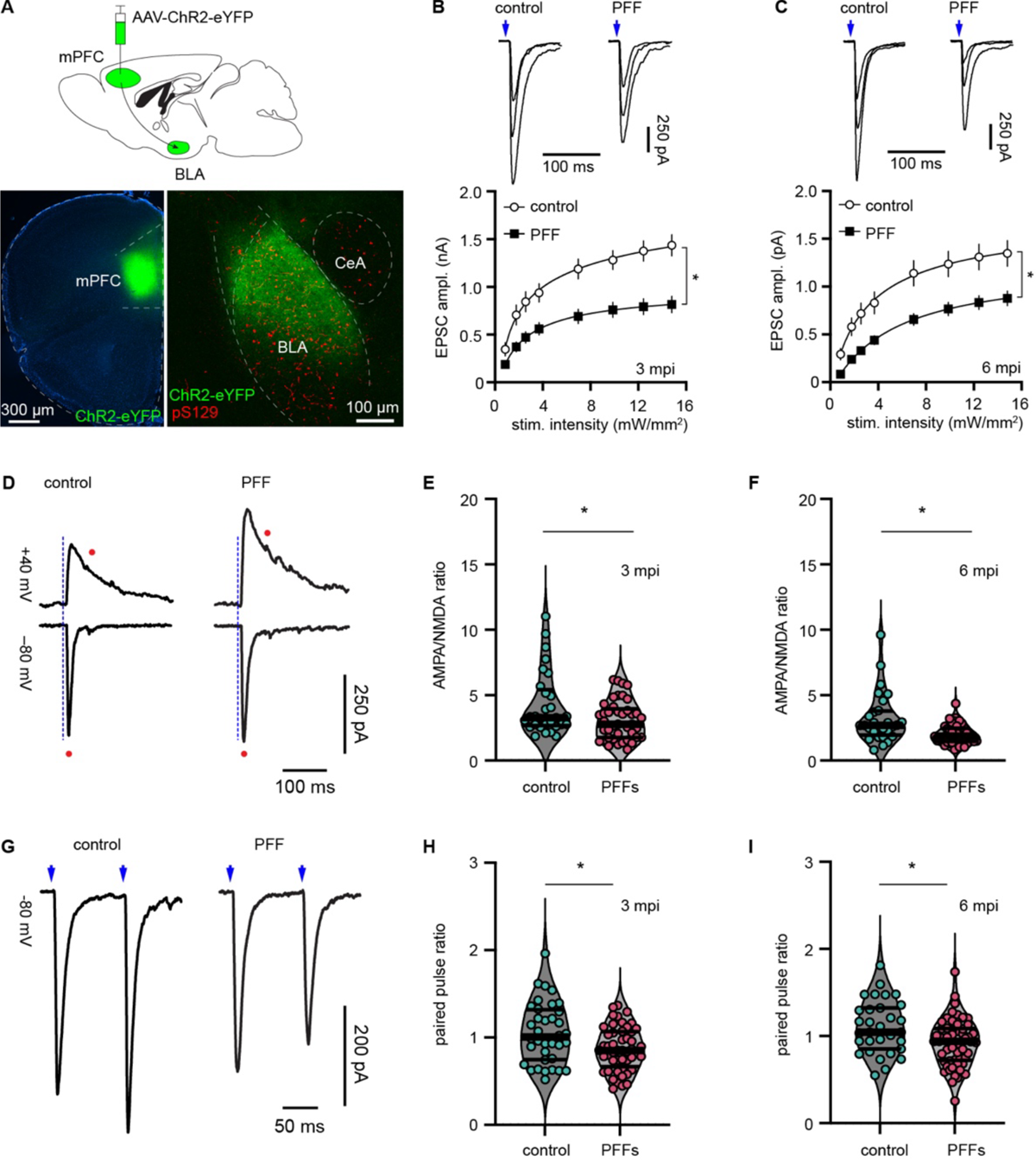
α-Syn pathology disrupts functional connectivity of the mPFC-BLA synapses. **A)** *Top*, diagram showing overall experimental design. *Bottom*, Representative AAV-ChR2-eYFP injection site in the mPFC (left) and the eYFP-labeled axonal terminal field in the BLA (right). **B-C**) Decreased amplitude of optogenetically evoked mPFC-BLA EPSCs in PFFs-injected mice relative to controls at both 3 (B) and 6 (C) mpi. *Top*) Representative traces of EPSCs evoked by optical stimulation intensities of 1.8, 3.6, and 12 mW/mm^2^. Blue arrows indicate when the optogenetic stimulation was delivered. *Bottom*) Summarized graphs showing reduced amplitude of mPFC-BLA ESPCs across a range of stimulation intensities in PFFs-injected mice relative to controls at both 3 and 6 mpi. N = 44 cells/7 mice at 3 mpi and 57 cells/7 mice at 6 mpi. Two-way ANOVA followed by Sidak multiple comparisons test. **D**) Representative traces of mPFC-BLA EPSCs recorded at −80 mV and +40 mV from both controls and PFFs-injected mice. Red dots indicate where the AMPA receptor- and NMDA receptor-mediated EPSCs were measured at −80 mV and +40 mV, respectively. Blue dashed lines indicate when optogenetic stimulation was delivered. **E-F**) Summarized result showing reduced AMPA/NMDA ratios at mPFC-BLA synapses in PFFs-injected mice relative to controls at 3 mpi (E, control = 3.3 [2.7, 5.4], n = 29 cells/4 mice, PFFs = 2.8 [1.8, 4.0], n = 41 cells/7 mice, p = 0.05) and 6 mpi (F, control = 2.7 [1.9, 3.8], n = 28 cells/4 mice, PFFs = 1.9 [1.5, 2.5], n = 46 cells/6 mice, p = 0.0006). **G**) Representative traces of mPFC-BLA EPSCs evoked by paired optogenetic stimulation pulses at −80 mV in both controls and PFFs-injected mice. Blue arrows indicate when the optogenetic stimulation was delivered. **H-I**) Summarized results showing reduced ratios of EPSC2/EPSC1 at the mPFC-BLA synapses in PFFs-injected mice relative to controls at both 3 mpi (H, control = 1 [0.75, 1.32], n = 36 cells/5 mice; PFFs = 0.85 [0.67, 1.1], n = 48 cells/7 mice, p = 0.016, MWU) and 6 mpi (I, control = 1.04 [0.85, 1.32], n = 31 cells/4 mice, PFFs = 0.94 [0.72, 1.09], n= 57 cells/7 mice, p = 0.008, MWU test).

The reduced connection strength of mPFC-BLA synapses was accompanied by a decreased ratio of amplitude of EPSCs mediated by AMPA receptors and NMDA receptors (i.e., AMPA/NMDAR ratio) in PFFs-injected mice relative to controls at both 3 and 6 mpi (Figure 4D-F). Moreover, the ratio of EPSC2/EPSC1 at the mPFC-BLA synapses evoked by paired optogenetic stimulation was also decreased in PFFs-injected mice relative to controls at both 3 and 6 mpi (Figure 4G-I), indicating an enhanced initial release probability of neurotransmitter at mPFC-BLA input. Together with the time course of synaptic marker changes (Figures 2 **and 3**), these data demonstrated that α-Syn pathology decreases the connection strength of cortico-BLA pathway at 3 mpi prior to a robust loss of cortico-BLA synapses at 6 mpi.

### Social interaction behavior is impaired by α-Syn pathology and rescued by chemogenetic activation of mPFC-BLA pathway

Given the functional disconnection between the mPFC and the BLA at 3 mpi (Figure 5), we next attempted to determine whether the impaired mPFC-BLA projection affects amygdala-dependent behavior. We performed behavioral tests using both controls and PFFs-injected mice at 3 mpi when the α-Syn pathology remained at peak levels but has not yet resulted in significant neuronal or synaptic losses in the BLA (Figures 2 **and 3**), making functional rescue still possible. We chose the 3-chamber sociability test, as the performance in this test is dependent on an intact amygdala microcircuit as well the interaction between the BLA and the mPFC [36, 39, 40]. Each test began with 10 min habituation to the behavior apparatus with two empty cages. During the habituation, both controls and PFF-injected mice spent comparable amount of time in the left and right chambers of the empty cage. This result indicates that neither controls nor PFFs-injected mice exhibited baseline preference of either chamber. Next, a stimulus mouse was placed into one empty cage (S1, Figure 5A), and the test mouse was then allowed to explore both chambers (session I in Figure 5A). Both controls and PFF-injected mice spent significantly more time with the never-before-met stimulus mouse S1 than that with the non-social empty cage (Figure 5A**, B, D**). This result suggests that PFFs-injected mice exhibit normal sociability relative to controls. Next, we studied social novelty preference of PFFs-injected mice. To do this, we added a second novel stimulus mouse 2 (S2) into the empty cage in the presence of the familiar stimulus S1 to evaluate the test mouse’s interest in the familiar social stimulus *versus* the novel one when both were present (see the session 2 in Figure 5A**, C**). While controls spent much more time with the novel stimulus (S2) relative to the familiar one (S1), PFFs-injected mice spent similar amount of time with either the familiar stimulus (S1) or the novel stimulus (S2) (Figure 5C**, E**).

**Figure 5.**
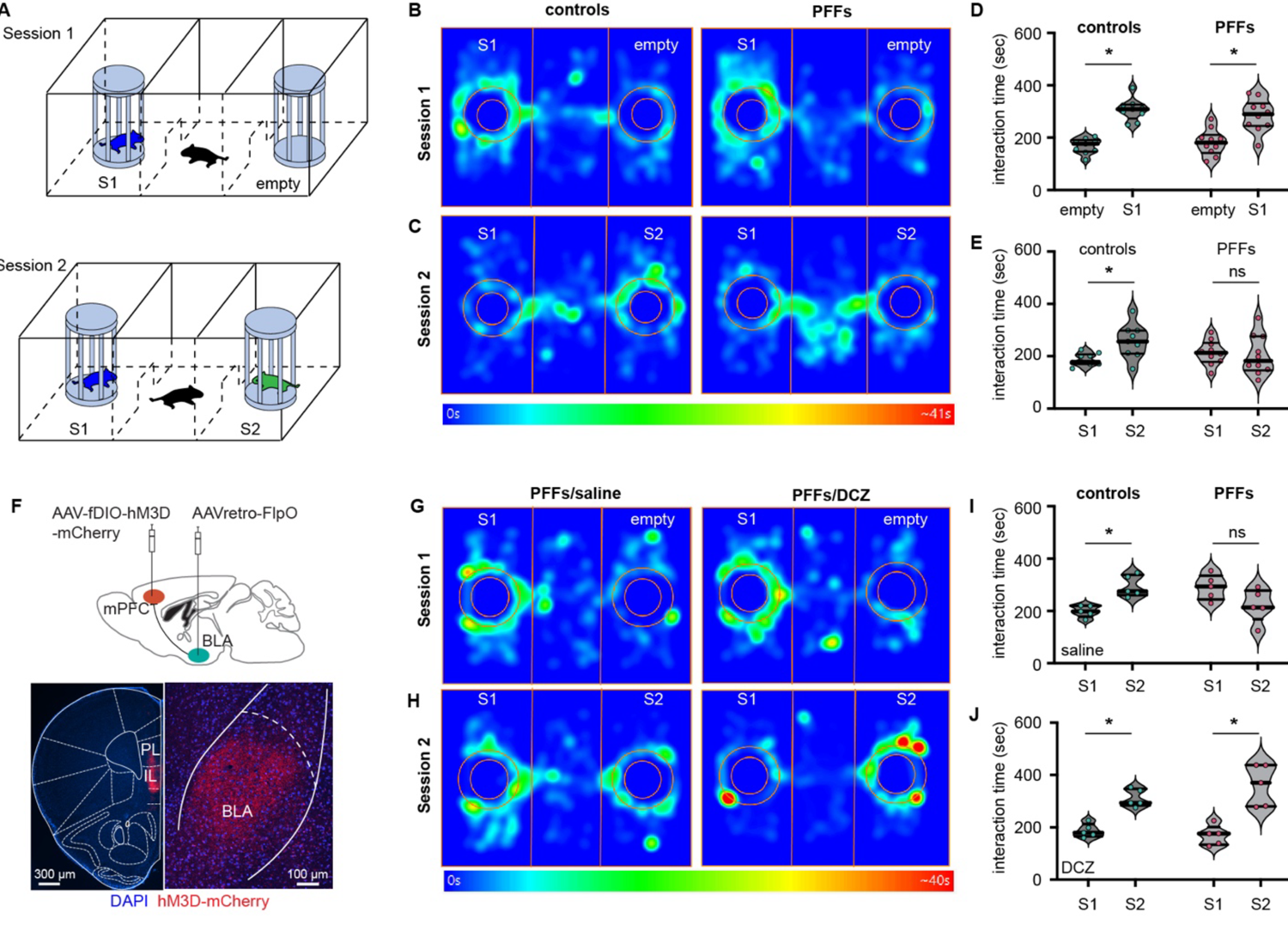
Social interaction behavior was impaired by α-Syn pathology and rescued by chemogenetic activation of mPFC-BLA pathway. **A)** Diagram showing overall behavioral experimental design. **B**) Representative heatmap plots showing social interaction behavior of controls and PFFs-injected mice in the Session 1 when S1 was paired with an empty cage for sociability. **C**) Representative heatmap plots showing social interaction behavior of controls and PFFs-injected mice in the Session 2 when familiar S1 was paired with a novel stimulus S2 for social novelty preference. **D-E**) Summarized graphs showing a normal sociability (D, controls empty = 177 [147, 191] sec, control S1 = 309 [273, 326] sec, n = 9 mice, p < 0.0001; PFFs empty = 181 [142, 211] sec, PFFs S1 = 290 [245, 331] sec, n = 10 mice, p = 0.001, MWU) but impaired social novelty preference behavior of PFFs-injected mice relative to controls (E, control S1 = 179 [170, 207] sec, control S2 = 256 [210, 298] sec, n= 9 mice, p = 0.01, MWU; PFFs S1 = 213 [179, 251] sec, PFFs S2 = 182 [146, 276] sec, n = 10 mice, p = 0.49, MWU). **F**) Representative image showing AAV-fDIO-hM3D(Gq)-mCherry infection site in the mPFC and the mCherry-labeled mFPC-BLA axon terminal field. **G**) Representative heatmap plots showing sociability of PFFs-injected mice receiving either saline or DCZ (DREADDs agonist) administration. **H**) Representative heatmap plots showing social novelty preference of PFFs-injected mice receiving either saline or DCZ (DREADDs agonist) administration. **I-J**) Summarized graphs showing that PFFs-injected mice receiving saline injections exhibit impairment in their social novelty preference behavior of (I, saline-injected control, S1 = 198 [178, 220] sec, S2 = 275 [260, 338] sec, n = 5 mice, p = 0.008; saline-injected PFFs, S1 = 294 [244, 335] sec, S2 = 214 [168, 278] sec, n =5 mice, p = 0.09, MWU), which was rescued by DREADDs agonist DCZ injection (J, DCZ-injected controls, S1 = 180 [166, 213] sec, S2 = 294 [284, 347] sec, n = 5 mice, p = 0.008; DCZ-injected PFFs, S1 = 176 [134, 201] sec, S2 = 371 [280, 438] sec, n = 5 mice, p = 0.008, MWU).

It is important to note that relative to controls, PFFs-injected mice at 3 mpi showed comparable general motor activity in locomotion test and rotarod test and similar levels of anxiety elevated plus maze as measured by the time spent in open arms (**Supplementary Figure 1**). These data were consistent with previous report [3, 25, 31] and suggested that PFFs-injected mice do not develop parkinsonian motor deficits or enhanced anxiety, which is important as such behaviors could confound the social interaction behavioral test. Taken together, these results suggested that PFFs-injected mice exhibited impaired social novelty preference and were not able to distinguish between familiar- and novel-stimuli.

Of particular note, while the mPFC-BLA projections showed significant functional impairment in PFFs-injected mice, there was no significant loss of their presynaptic axonal terminals at 3 mpi. Given the critical role of mPFC inputs to the BLA in regulating social interaction, we sought to determine whether the deficient social novelty preference behavior could be rescued via activation of mPFC-BLA connections in PFFs-injected mice. To this end, we injected retrograde AAV-FlpO into the BLA and AAV-fDIO-hM3D(Gq)-mCherry into the mPFC (Figure 5F). This intersectional approach allows us to selectively stimulate mPFC-BLA synaptic neurotransmission using a chemogenetic approach. Three weeks after viral infections, PFFs-injected mice were subject to intraperitoneal injections of saline or deschlorozapine (DCZ) followed by social novelty preference behavior test. After saline injection, while controls showed normal social novelty preference, PFFs-injected mice were not able to distinguish novel-*versus* familiar-stimulus mice (i.e., S2 *vs* S1, Figure 5H**, I**), consistent with our data from previous cohort of animals (Figure 5C, E). In contrast, when PFFs-injected mice were subject to social novelty preference test after DCZ injection, they showed strong preference to S2 *versus* S1 (Figure 5J), suggesting that their impaired social novelty preference has been rescued by chemogenetic stimulation of mPFC-BLA connection.

Altogether, the above data suggest that the social novelty preference was disrupted by α-Syn pathology in PFFs-injected mice at 3 mpi. Importantly, such disruption correlated with impaired mPFC-BLA synaptic connection, which was then rescued by chemogenetic stimulation of the mPFC-BLA connection in the PFFs-injected mice.

## Discussion

In the present study, we performed longitudinal analyses of anatomical and physiological changes associated with the formation of α-Syn aggregation, and provided novel insight into how α-Syn aggregation impairs the connectivity and function of amygdala microcircuits. While most research in the field has focused on the processes that initiate, facilitate, and/or exacerbate α-Syn pathology formation and propagation, the present work highlights the less explored, yet important, circuitry dysfunction associated with α-Syn aggregation that can underlie some of the symptoms manifested by individuals afflicted by synucleinopathies. These studies are of clinical relevance as strategies to prevent worsening of α-Syn pathology and to repair the damaged circuits could make the current treatments for diseases like PD and DLB more effective.

### Synaptic dysfunction occurs earlier than neurodegeneration in the BLA

We demonstrated that α-Syn aggregation compromises synaptic function before structural changes occur, including the loss of presynaptic axon terminal markers. In the context of our recent work, even though functional deficits of cortico-BLA synapse may occur as early as 1 mpi [34], loss of cortico-BLA presynaptic marker vGluT1 was only detected at the 6 mpi timepoint (Figure 2). Moreover, BLA neurons are also prone to accumulating α-Syn aggregates, which perhaps results in the loss of spines at 3 mpi (Figure 4) and their degeneration at much later stages, i.e., 12 mpi. These observations are consistent with reports from the nigrostriatal system in parkinsonian animals, showing deficits in dopamine release and degeneration of axons prior to severe degeneration of dopaminergic neurons [4, 6, 17, 41, 42].

It is important to note that while BLA neurons exhibited loss of dendritic spines, the initial release probability at cortico-BLA synapses increased at 3 and 6 mpi (Figure 4). This observation is consistent with earlier findings from cultured hippocampal neurons, in which the loss of postsynaptic spines and a paradoxical increase of presynaptic glutamate release were reported [43]. Froula *et al* suggested that the enhanced presynaptic release probability might be due to an increased size of readily releasable pool of neurotransmitters.

### Synaptic dysfunction is sufficient to drive the manifestation of behavioral phenotypes

At the early stage of α-Syn based models, functional alterations of nigrostriatal pathway have been reported to underlie the manifestation of parkinsonian motor deficits [4, 17, 44, 45]. In the present study, we found that, prior to the loss of BLA neurons, mPFC-BLA synapses exhibited functional impairment in PFFs-injected mice at 3 mpi, which temporally correlated with the manifestation of defected social interaction behavior (Figures 4 **and 5**). Interestingly, the impaired social novelty preference behavior was rescued by chemogenetic stimulation of the defected mPFC-BLA pathway. Together, these findings support the hypothesis that synaptic dysfunction is sufficient to drive the manifestation of behavioral phenotypes in synucleinopathies. The intrastriatal PFFs-injected mice exhibited very specific deficits in emotion related behavior. For example, while they exhibited deficits in social novelty preference (Figure 5) and social dominance behavior [31], no change in their anxiety levels in the elevated plus maze test were detected [25, 31]. Though compelling evidence suggests that both social behavior and anxiety are tightly regulated by the amygdala, it is plausible that performance in the elevated plus maze test may involves distinct amygdala subcircuits from those activated during the 3-chamber social interaction test [46, 47].

### Cell type-specific α-Syn expression and its implications in circuit dysfunction

In a complex circuit, not all afferents within the same neurochemical category are equally affected by α-Syn aggregation. For example, vGluT2^+^ glutamatergic axons arising from the thalamus are more resilient relative to vGluT1^+^ axons in the BLA (Figure 2). This observation is largely consistent with lower *Snca* mRNA levels in thalamic neurons as well as α-Syn expression in their axon terminals [34, 48]. Moreover, evidence in the literature consistently reports that Lewy body-like pS129^+^ somatic aggregates are more abundant in cerebral cortical regions relative to thalamic regions in intrastriatal PFFs models [3, 25, 26, 34, 49], suggesting that the cerebral cortex and thalamus are affected differently by α-Syn pathology in diseased state. Of relevance to the present work, it is known that cooperative activity and plasticity of cortical and thalamic inputs are essential to normal amygdala function in the formation and long-term storage of emotion memory [50, 51]. It remains to be determined how the disrupted balance or cooperation of cortical and thalamic inputs to the BLA might impair emotion encoding and regulation by the amygdala circuits. Remarkably, human imaging studies reported that PD individuals diagnosed with depression showed a reduced activation of the amygdala in the presence of aversive stimulations as well as a disconnected cortical (but not the thalamic) inputs to the amygdala [27, 30]. Our findings of weakened cortical, but not thalamic inputs, to the BLA might be of relevance to this clinical observation in PD individuals exhibiting neuropsychiatric symptoms.

The complicated interaction and impact of α-Syn pathology on circuit function can also partially explain why emotion dysregulation, including anxiety and depression, can occur at any stages of the disease in PD or DLB individuals (e.g., even before the onset of motor symptoms), which might vary depending on where the α-Syn pathology starts and how it can affect specific circuits. While the present studies focus on the impact of α-Syn pathology on the amygdala function and related behavior, we must acknowledge that degeneration of key neuromodulators and their inputs to the amygdala (e.g., dopaminergic and noradrenergic systems) also play key roles in the development of neuropsychiatric symptoms in PD and DLB.

## Conclusion

We presented a combination of anatomical and physiological evidence to demonstrate a series of circuitry events occurred in the amygdala circuits associated with α-Syn pathology in mouse brain. Our findings not only support several well-accepted concepts in the field (e.g., synaptic deficits prior to neurodegeneration), but also highlight the fact that we need to consider circuit complexity before we can gain a thorough understanding on the contribution of α-Syn to brain function in health and in diseased states.

## List of abbreviations

BLA: basolateral amygdala

α-Syn: α-synuclein

PD: Parkinson’s disease

DLB: Dementia with Lewy bodies

SNc: Substantia nigra pars compacta

PFFs: α-Syn preformed fibrils (PFFs).

EPSCs: excitatory postsynaptic currents

mPFC: medial prefrontal cortex

DREADDs: designer receptors exclusively activated by designer drugs IL-mPFC infralimbic subregion of mPFC

aCSF: artificial cerebrospinal fluid

Rs: series resistance

PFA: paraformaldehyde

ANOVA: analysis of variance

pS129^+^: phospho-Ser129

LN: Lewy neurite

LB: Lewy body

mpi: months-post-injection

vGluT1: vesicular glutamate transporter 1

vGluT2: vesicular glutamate transporter 2

SNCA: Alpha Synuclein

DCZ: deschloroclozapine

## Acknowledgement

Experimental work was conducted at Van Andel Research Institute. Data analysis was partially performed at Van Andel Research Institute and completed at Georgetown University Medical Center. The authors thank Van Andel Institute vivarium technician team for animal husbandry, and Ms. Rachel Dvorak for initial technical assistance in animal surgery. The authors thank Dr. Qiang Zhu at Van Andel Institute for general support in grant administration.

---

**Supplementary Figure 1.**
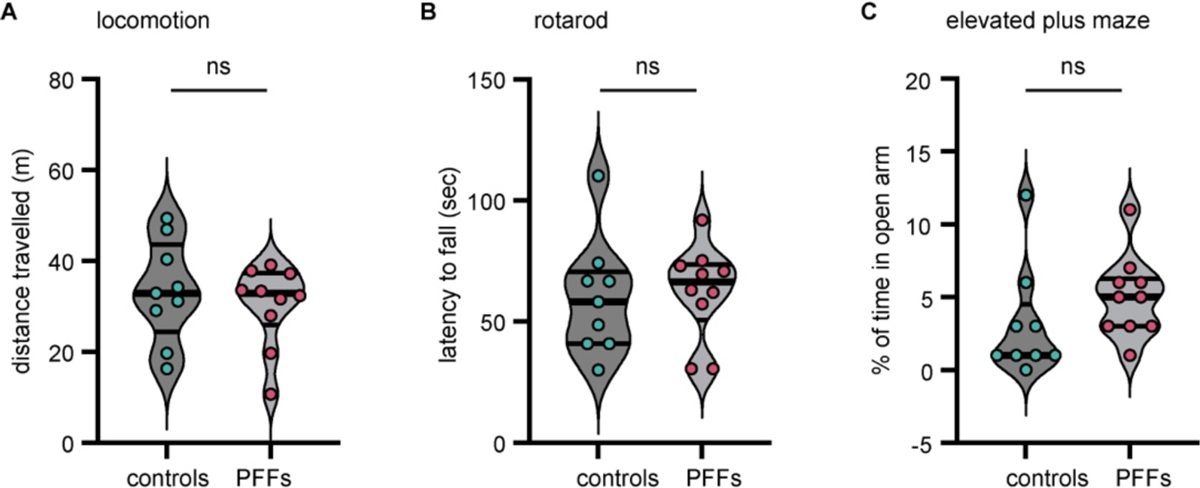
PFFs-injected mice showed normal motor function and anxiety level at 3 mpi. **A-C**) Summarized graphs showing no difference between controls and PFFs-injected mice in the locomotion test (A, distance travelled, control = 33 [24, 44] m, n = 9 mice, PFFs = 33 [26, 37], n = 10 mice, p = 0.7, MWU), the rotarod test (B, latency to fall, control = 58 [41, 71] sec, n = 9 mice, PFFs = 66 [51, 74] sec, n = 10 mice, p = 0.5, MWU) and the elevated plus maze (C, % of time in open arm, control = 1 [1, 4.5]%, n= 9 mice, PFFs = 5 [3, 6]%, n = 10 mice, p = 0.08, MWU).

